# Biases in hand perception are driven by somatosensory computations, not a distorted hand model

**DOI:** 10.1101/2023.12.06.570330

**Authors:** Valeria C. Peviani, Luke E. Miller, W. Pieter Medendorp

## Abstract

To sense and interact with objects in the environment, we effortlessly configure our fingertips at desired locations. It is therefore reasonable to assume the underlying control mechanisms rely on accurate knowledge about the structure and spatial dimensions of our hand and fingers. This intuition, however, is challenged by years of research showing drastic biases in the perception of finger geometry.^1–5^ This perceptual bias has been taken as evidence that the brain’s internal representation of the body’s geometry is distorted,^6^ leading to an apparent paradox with the skillfulness of our actions.^7^ Here, we propose an alternative explanation of the biases in hand perception—They are the result of the Bayesian integration of noisy, but unbiased somatosensory signals about finger geometry and posture. To address this hypothesis, we combined Bayesian reverse-engineering with behavioral experimentation on joint and fingertip localization of the index finger. We modelled the Bayesian integration either in sensory or in space-based coordinates, showing that the latter model variant led to biases in finger perception despite *accurate* representation of finger length. Behavioral measures of joint and fingertip localization responses showed similar biases, which were well-fitted by the space-based but not the sensory-based model variant. The space-based model variant also outperformed a distorted-hand model with built-in geometric biases. In total, our results suggest that perceptual distortions of finger geometry do not reflect a distorted hand model but originate from near-optimal Bayesian inference on somatosensory signals.

## RESULTS

One of the most surprising neuroscientific findings over the last two decades is that the spatial perception of our body appears to be highly distorted, even for body parts that require fine-grained motor control, such as the hand and fingers.^1^ A widespread assumption is that the perceptual biases measured in these experiments *directly* reflect a distorted internal representation of body geometry ^6,8^ (but see ^9,10^). Taking this inference at face value for the hand leads to what has been termed the ‘hand paradox’:^7^ How are distorted hand representations compatible with the observed skillfulness of our manual actions?

Here, we address this paradox. In contrast to the standard interpretation, we propose that the observed perceptual biases do not reflect a distorted hand model; rather, these biases arise from probabilistic computations that perform near-optimal Bayesian inference on somatosensory signals. We took a two-pronged approach to investigate whether this optimal probabilistic processing can explain the perceptual biases observed in a finger mapping task: First, we formalized and modelled the optimal computations that may underlie the transformation of somatosensory signals about finger configuration into its spatial dimensions (**Figure 1**). This allowed us to obtain quantitative predictions pertaining to spatial errors in finger mapping. We then tested these predictions using a novel VR-based finger mapping task, combined with fitting our computational models to the observed responses (**Figure 2**).

**Figure 1.**
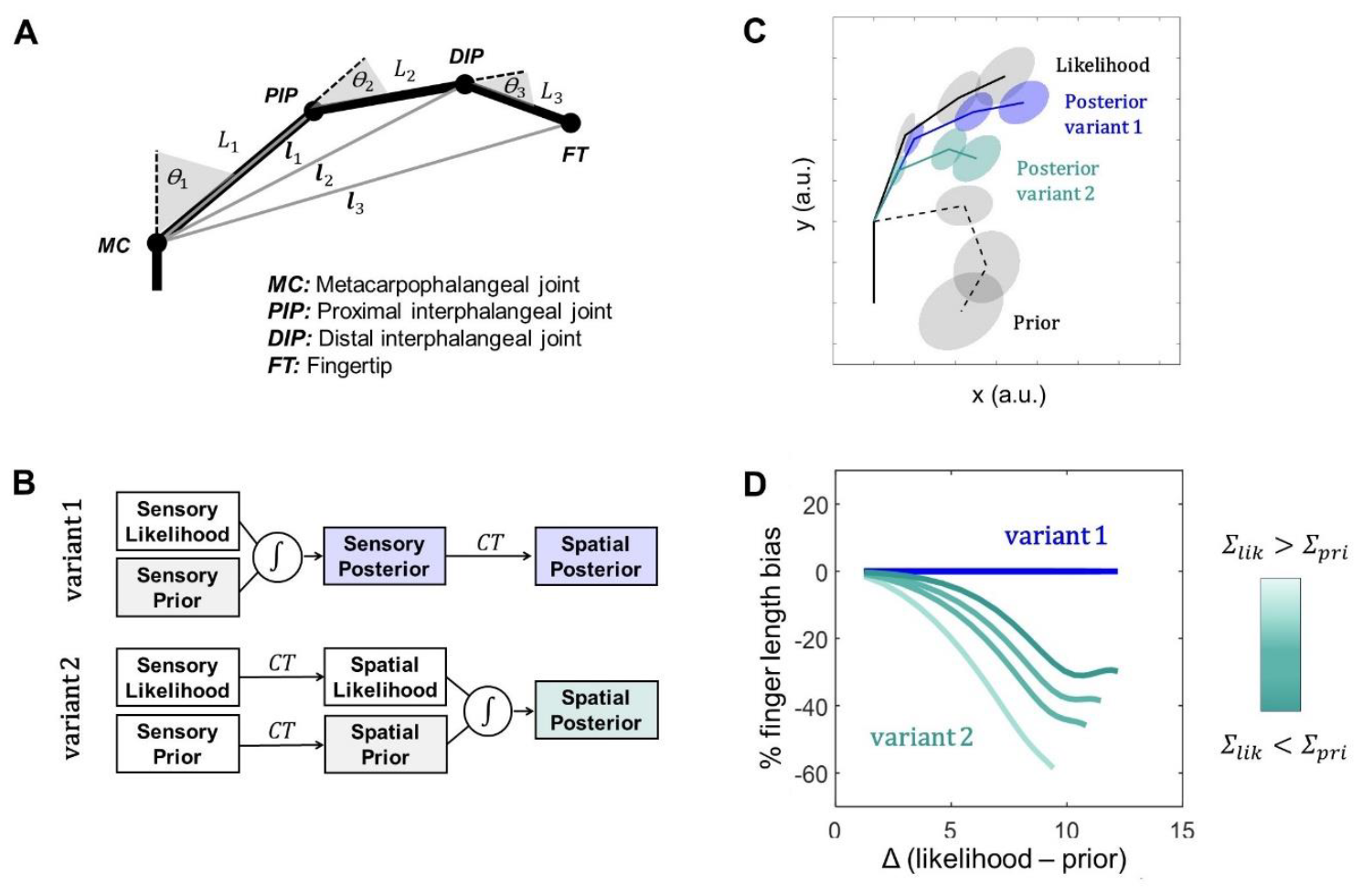
Bayesian model of finger perception. **A**. Geometric model of the finger as a kinematic chain. The chain allows one-axis rotations. The three segments (*L*_1_, *L*_2_, *L*_3_) are interconnected by three joints (*θ*_1_, *θ*_2_, *θ*_3_), corresponding to the MC, PIP and DIP. The spatial positions of PIP, DIP and FT are defined as 2D vectors ***l***_**1**_, ***l***_**2**_, ***l***_**3**_, each with respect to MC. **B**. Schematic representation of the Bayesian model of finger perception. Variant 1: Bayesian integration (∫) of signals encoding finger joint angles and length, before the sensory-to-space transformation (*CT*). Variant 2: Bayesian integration occurs in Cartesian coordinates, after the sensory-to-space transformation. **C**. Example of model predictions: both variants plotted in Cartesian coordinates (likelihood, solid gray; prior, dashed gray; posterior variant 1, blue; posterior variant 2, green). Variant 1 predicts biases in joint angle, not phalanx length. Variant 2 predicts biases in both, as if the overall length of the finger is underestimated. **D**. Finger length bias plotted as a function of the Mahalanobis distance between likelihood and prior, Δ*(likelihood-prior)*, which quantifies their distance in statistical terms. Variant 2 predicts that this bias increases with this distance (see also **Figure S1, SI**).

**Figure 2.**
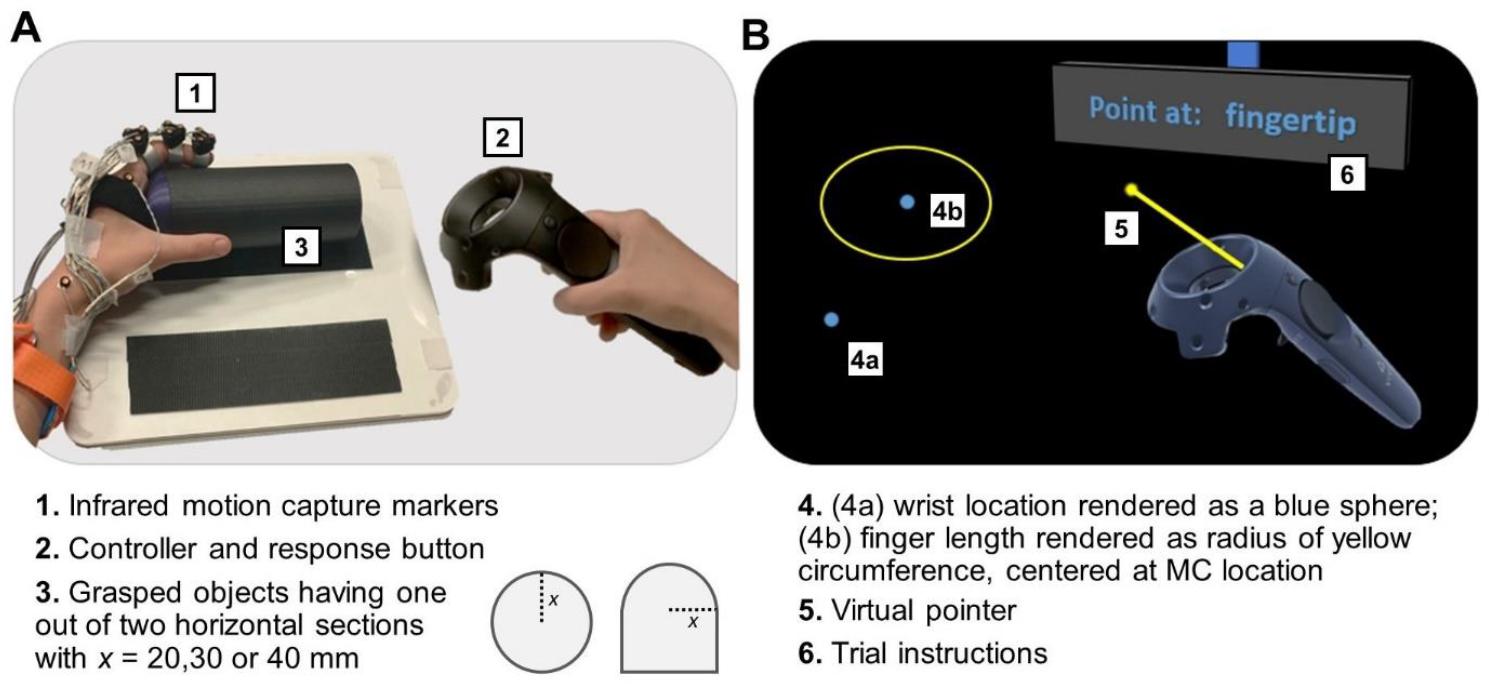
Experimental task and stimuli. **A**. Participants wore a VR headset, while their head was supported by a chin rest. In each block of 100 trials (six blocks in total), participants adopted a unique figure configuration by firmly grasping an object of variable size and shape with a power grip (see also **Figure S2, SI**). The positions of left wrist, left index metacarpophalangeal (MC), proximal interphalangeal (PIP), distal interphalangeal (DIP) joints and fingertip (FT) were recorded using infrared motion tracking. **B**. The VR environment presented a dark 3D space in which experimental instructions were displayed on a screen. Blue spheres (diameter 5 mm) indicated the physical locations of participants’ left wrist and MC. Information on total index finger length was conveyed by rendering a yellow circle centered on MC. Participants used a hand-held virtual pointer to indicate the perceived locations of PIP, DIP or FT.

### A Bayesian model of finger perception

The structure of a finger is characterized by its joints, whose angles are derived from proprioceptive senses,^11^ and by phalanges, whose lengths can only be estimated indirectly from other somatosensory signals; for example, from changes in muscular torque, skin stretch or changes in motor output.^12–14^ We developed a model of how these signals about finger geometry are transformed into a percept of where the finger is in space. To do so, we conceptualized the finger as a kinematic chain, where each phalanx having length *L* is linked with the previous phalanx through their interphalangeal joint, having angle *θ*. As illustrated in **Figure 1A**, this information allows one to derive the Cartesian positions of the joints and the fingertip relative to the metacarpophalangeal joint, following:

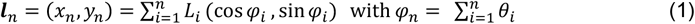

where ***l***_*n*_ is the two-dimensional location of the proximal (***l***_1_) or distal (***l***_2_) interphalangeal joint or the fingertip (***l***_3_). However, this formulation is not sufficient to specify the underlying computations used by the nervous system. The brain does not represent the angles, lengths, and locations as individual point estimates but rather as probability distributions.^15^ This is because the activity within the nervous system is intrinsically noisy, from transduction of signals to network interactions.^16^ To deal with this uncertainty, the brain must rely on probabilistic processing, as formalized by Bayesian decision theory.^17,18^ For perception, the Bayesian framework states that the observer forms an estimate (the posterior) about the most probable state of the body and the world by integrating noisy sensory signals (the likelihood) with prior beliefs (the prior), following Bayes’ rule. Importantly for our purposes, this computation yields an increase in perceptual precision at the expense of a bias in perception.

We can therefore re-formalize the geometric model of the finger in Eq. (1) in probabilistic terms (**Figure 1B**). The spatial percept of the finger (posterior) is constructed by integrating the somatosensory measurements (likelihood) about the finger configuration with prior information about its most probable configuration (prior), possibly inferred from an accumulated history of the finger’s previous configurations.^18^ Since the size of our fingers remain stable in the short-term, it is reasonable to assume that prior information on segment length *L* is accurate, i.e., centered on true values.

For each ***l***_*n*_, we specify the likelihood and the prior as vectors with their respective means (***μ***_*lik*_ and ***μ***_*pr*_) and diagonal variance-covariance matrices (*∑*_*lik*_ and *∑*_*pr*_). Following Bayes’ rule, the posterior is obtained from the product of the likelihood and prior. In matrix-form, this is formalized as follows:

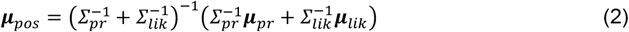

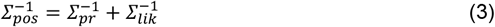

where Eq. (2) is the computation of the posterior mean (***μ***_*pos*_) and Eq. (3) is the computation of the posterior variance-covariance matrix (*∑*_*pos*_), assuming that the Bayesian integration is performed optimally. Note that Eq. (3) represents *∑*_*pos*_ on a single trial. However, in an experiment, we measure the response distribution (*∑*_*r*_) across trials.^19,20^ It can be shown that *∑*_*r*_ = *W * ∑*_*lik*_ *\* W*^*T*^, where 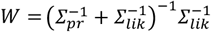, see **STAR Methods**.

While Bayesian integration could occur at different sensory stages,^21–24^ here we consider two distinct model variants that formalize how the sensory-to-space transformation (Eq. 1) and Bayesian Integration (Eqns. 2 and 3) are implemented in the somatosensory processing hierarchy (**Figure 1C**).

In **model variant 1** (**Figure 1B**, top; **Figure 1C**, blue), Bayesian integration occurs *before* the sensory-to-space transformation. That is, the likelihood and the prior in Eqns. (2-3) are defined in sensory coordinates, representing the involved joint angles (*θ*_1_, …, *θ*_*n*_) and phalanx lengths (*L*_1_, …, *L*_*n*_). The likelihood is specified as a Gaussian distribution with mean ***μ***_*lik*_ and diagonal variance-covariance matrix *∑*_*lik*_, where all off-diagonal are zero (i.e., assuming uncorrelated signals). The prior is likewise specified as Gaussian with mean ***μ***_*pr*_ and variance-covariance matrix *∑*_*pr*_; however, unlike the likelihood, we did allow covariance between subsequent joints. Following Bayesian integration (Eqns. 2-3), the posteriors of the joint angles and segment lengths are then transformed into space-based posteriors of the joints or fingertip, involving the Jacobian transformation as derived from Eq. (1), see **STAR Methods**.

In **model variant 2** (**Figure 1B**, bottom; **Figure 1C**, green), Bayesian integration occurs *after* the sensory-to-space transformation. In this variant, the sensory likelihood distributions are first transformed into space-based likelihoods of joint and fingertip position (i.e., ***μ***_*lik*_ and *∑*_*lik*_ specified in Cartesian coordinates). These are then integrated with the transformed spatial prior (***μ***_*pr*_, with *∑*_*pr*_) as in Eqns. (2-3) to obtain the space-based posterior of the joints and fingertip positions.

### Near-optimal computations predict perceptual biases in finger perception

We next identified the differences in the quantitative predictions made by each model variant. Our goal here was two-fold. First, we aimed to distinguish specific testable predictions made by each model. Second, and relatedly, we aimed to pinpoint when—if ever—these computations would lead to distorted perception of joint and fingertip locations. To do so, we simulated the above computations across a variety of model parameters and finger postures.

**Figure 1C** illustrates the computational outcomes of each model in a Cartesian reference frame. In both model variants the posterior distribution is biased relative to the actual joint position. Crucially, only in variant 2 do these biases affect the perceived geometry of the finger, as inferred from the position estimates of each joint/tip. Interestingly, these finger length distortions were a ubiquitous consequence of Bayesian integration occurring after the sensory-to-space transformation. This is despite the fact that the underlying likelihood and prior accurately reflected finger size.

Simulations with model variant 2 (**Figure S1, Supplemental Information, SI**) further revealed that the magnitude of finger length misestimation varied as a function of posture. **Figure 1D** illustrates how the percentage of total finger length misestimation varies as a function of the Mahalanobis distance between the spatial likelihood and the spatial prior. As this distance increases, only model variant 2 predicts the emergence of length misestimation. Hence, the posture-dependence of the inferred finger length is the key difference between the predictions of the two model variants. Crucially, this posture-dependent misestimation would not be predicted if perceptual biases were only a consequence of distorted representation of finger length (see below).

### Probing model predictions in a novel VR finger-mapping paradigm

To test the model’s predictions, we designed a novel VR-based finger mapping task (**Figure 2**, and **STAR Methods**). In each block (six in total), participants adopted a unique figure configuration by firmly grasping an object of variable size and shape (**Figure 2A** and **Figure S2, SI**). On each trial, participants used a hand-held virtual pointer to localize the position of one of three landmarks (i.e., both interphalangeal joints and fingertip) on their unseen index finger (**Figure 2B**). Participants performed 100 trials in each block (600 in total). We used a reverse-engineering approach to identify which model (**Figure 1B**) provided the best fit to each participant’s trial-by-trial position estimates (**STAR Methods**).

The results of our experiment matched the predictions from model variant 2. **Figure 3** illustrates the data of four representative participants, showing that the perceived location of finger joints and tip differed from their actual locations (**Figure 3A, Figure S4, SI**). This finding is consistent with the notion that Bayesian priors bias perception. Crucially, the observed biases took on two patterns: First, the perceived angle of the finger deviated from the actual angle, suggesting the role of a prior over joint angles in perception; Second, the perceived finger length—as derived from the position estimates—differed from the actual finger length, with an average underestimation of about 20% (mean±sem: 19.5 ± 4.4%). Our findings suggest that perceptual biases may arise from Bayesian computations occurring in Cartesian space, after the sensory signals are transformed into spatial coordinates.

**Figure 3.**
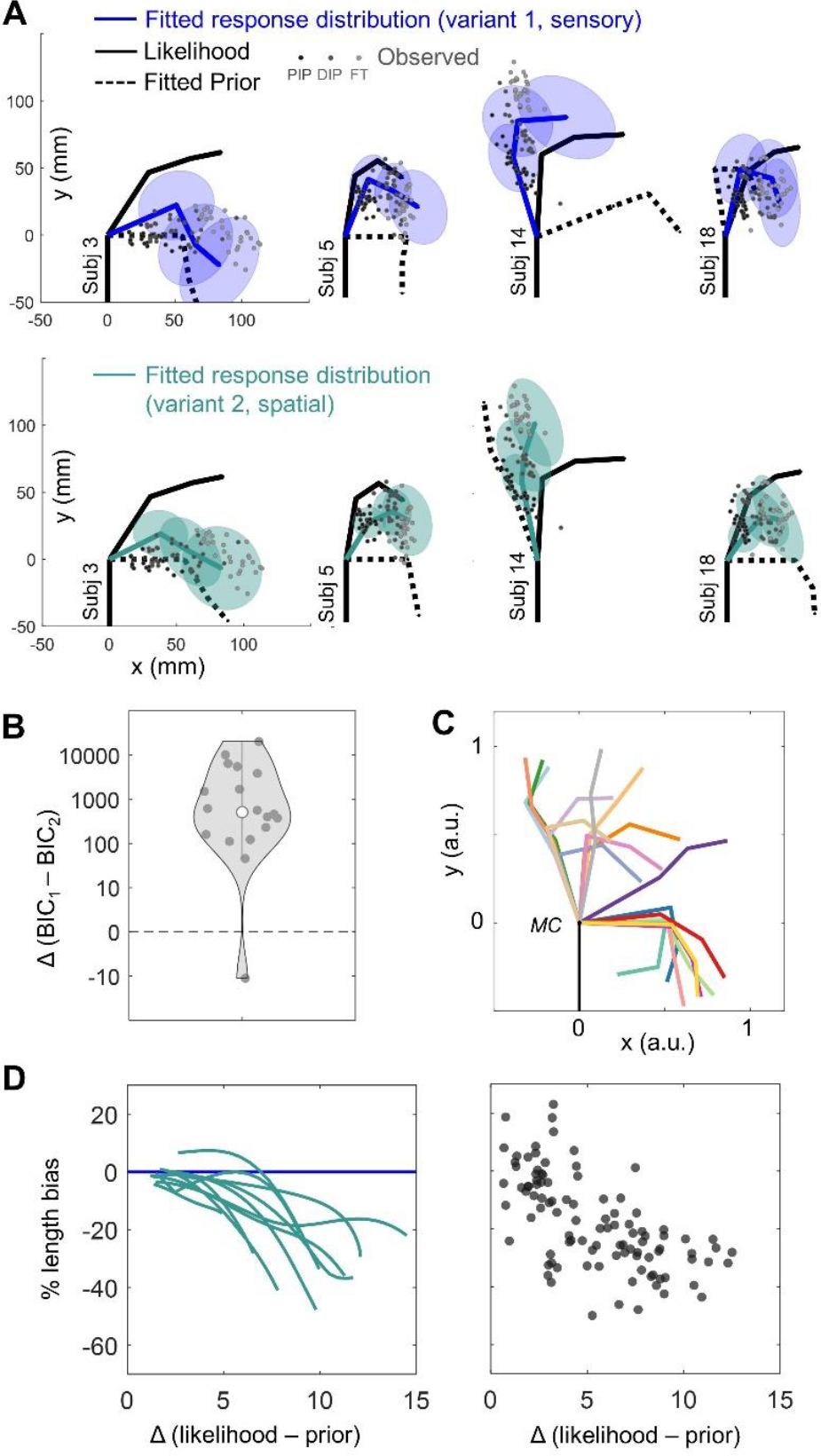
Near-optimal computations underlie biases in finger perception. **A**. Data of four representative participants for one of six finger configurations (in black). Localization responses of PIP, DIP and FT are represented as gray dots and their means are connected by gray lines. Model fits are shown as the covariance matrix of the response distribution for these locations (variant 1, in blue; variant 2, in green). The fitted prior mean is represented as a dashed line. See also **Figure S4, SI. B**. Model variant 2 outperformed variant 1 for 17 out of 18 participants, with a median (±iqr) BIC difference of 512 ± 369. **C**. Fitted prior means (variant 2) for all participants are plotted in normalized Cartesian units. **D**. Bayesian integration predicts posture-dependent biases in finger perception. We simulated spatial posteriors based on bestfit model variants of each subject for several finger configurations ranging from fully extended to flexed. Based on these posteriors, we plot the bias in finger length as a function of the Mahalanobis distance between likelihood and prior, *Δ(likelihood-prior)*. Predictions are represented as lines in the left panel, observations are represented as solid dots in the right panel. The predictions of a distorted hand model would be the same as model variant 1, but shifted at the bias parameter value (e.g., -20%).

Model fitting confirmed these initial observations. The fits of each model variant to the response distribution of representative participants can be seen in **Figure 3A** and **Figure S4, SI**. For almost every participant (17 out of 18), integration after the sensory-to-space transformation (model variant 2) best explained the behavioral observations, with consistently lower BICs compared to model variant 1. The median (±iqr) BICs were 8615 ± 4983 for variant 1 and 8105 ± 624 for variant 2, with a median difference of 512 ± 369, corresponding to a Bayes Factor (BF) of BF_v2,v1_ = 151e+109 (**Figure 3B**). Unlike model variant 1, model variant 2 predicts posture-dependent finger length misestimation biases. These predictions were correlated with the observed behavior (Pearson’s *r* = 0.37, *p*<.001): the larger the spatial discrepancy between the likelihood and prior, the greater the magnitude of finger length underestimation (**Figure 3D**). Furthermore, prior posture varied across participants (**Figure 3C, Table S2, SI**), leading to different patterns of length misestimation that peaked at different finger postures in different individuals.

Even though Bayesian model variant 2 provided a good description of our data, it is possible that the observations are better explained by the traditional view, where biases in finger perception stem from a biased hand representation. To explore this possibility, we formulated a probabilistic model of finger representation (distorted hand model) with built-in biases on phalanx length and joint angles (**STAR Methods**). We found that variant 2 always outperformed this model with a median (±iqr) BIC difference of 562 ± 605, corresponding to a BF_bias,v2_ = 159e+120. Indeed, the distorted hand model was unable to capture the posture-dependent nature of finger length misestimation that was pervasive in our participant’s judgments.

One potential methodological limitation of this experiment may be that participants during the VR task were presented with a circumference centered on their left MC (**Figure 2B**), which was designed to minimize spatial misestimation due to the virtual environment. It is possible that this cue itself may have induced a bias in position estimates made by participants. In contrast to this possibility, follow-up analyses suggested that it is unlikely that finger underestimation─and specifically its posture dependency─was due to this methodological choice; We discuss this point in detail in the **STAR Methods** and **Figure S5, SI**.

In total, our findings bring to light two key features of the computation used by the brain to transform somatosensory inputs into the perception of the body in space. First, estimating the finger’s position in space results from optimal integration of transformed sensory inputs with stored priors. Second, optimal integration between sensory and stored signals occurs between estimates of finger posture in a space-based, Cartesian reference frame. Together, the features of this computation lead to the consistently observed distortions in finger length perception, as inferred from position estimates. We discuss the implications of these results in the following section.

## DISCUSSION

Previous empirical observations suggest that postural priors influence proprioceptive and tactile processing. For example, during temporary limb anesthesia, limb perception systematically changes towards a semi-flexed posture.^25,26^ Placing the fingers or upper limbs in unusual postures leads to less efficient tactile processing.^27–34^ Going beyond these observations, our work is the first to quantitatively define and empirically verify the influence of a postural prior on the perception of the body in space. More crucially, our findings constrain the nature of this prior, suggesting that it is encoded in spatial (Cartesian) and not sensory coordinates (**Figure 1C**). A consequence of this computational scheme is that Bayesian integration occurs only after sensory signals are themselves transformed into a spatial estimate of finger location.

The configuration of the Cartesian priors varied across participants, though they tended to cluster near a more extended or flexed finger posture (**Figure 3C**). These extremes are common resting hand postures, suggesting that the Cartesian prior may reflect the accumulated history of previous finger configurations, built upon sensorimotor^35–37^ and visual^38^ feedback. Investigating the statistics of natural everyday finger postures would be one way address the origin of these priors.

There are important functional reasons why the brain would perform somatosensory inference in Cartesian spatial coordinates. For example, doing so would aid in multisensory integration (e.g., vision and proprioception), where sensory signals must be transformed from their native sensor-based co-ordinates into a shared spatial frame of reference. Bayesian multisensory integration is indeed a basis for body perception,^20^ the bodily self,^39–41^ and sensorimotor control.^42^ We propose that the model of the present study reflects the first step in these processes, transforming signals into a reference frame that could be shared amongst different sensory modalities, including the construction of peripersonal space.^43–46^

A major area of inquiry over the last decade has been the origin of distortions in hand perception. According to a recent conceptual model of body representation,^8^ postural priors are not sufficient to explain metric distortions in hand perception. Instead, perceptual distortions are the direct consequence of integrating somatosensory inputs with a distorted body representation (i.e., a body model). Both our simulations and empirical findings stand in contrast to this claim. Indeed, a Bayesian model with completely undistorted signals provided a better fit to every participant’s data than a model of finger representation with built-in distortions. Our results thus suggest that distortions in finger length perception are due to the nature of the computation; Specifically, spatial finger perception is the result of Bayesian integration between a spatial likelihood and prior (i.e., model variant 2) that each *accurately* encode finger length. One unique piece of evidence in favor of the present proposal is the posture-dependence on finger length misestimation (**Figure 3** and **Figure S4, SI**). These findings reconcile the notion of perceptual biases in finger perception with optimal sensorimotor processing, as all signals used for control would be unbiased.

One question is whether the computations we describe can also explain the results found in the classic hand-mapping task,^1^ which typically reports an underestimation of finger length of about 28%.^5^ In this task, participants indicate the position of hand landmarks on an occluding board position over their hand, which is lying flat on a table with fingers fully extended. Unlike our VR-based task, this approach would be insensitive to a posterior estimate of finger posture that would be curved in 3D space. We therefore simulated response distributions based on the parameters recovered for each participant (variant 2) for such a flat finger configuration. Average (±sem) underestimation of finger length as inferred from simulated position estimates was 31.1 ± 6.9%, which aligns very well with previous hand-mapping results.

Neurophysiological studies suggest that the computations identified in our study may occur as early as primary somatosensory cortex (S1). Previous research suggests that most neurons in each subdivision, including areas 3a and 3b, respond to cutaneous *and* proprioceptive signals, showing multimodal interactions among information encoded in different coordinates.^47^ Indeed, multimodal neurons in Area 2 are known to encode the posture of the forelimb in Cartesian coordinates.^48^ Furthermore, two recent human fMRI studies reported that activity in S1 correlates with the spatial features of the hand and biases in hand perception, as if it embeds a spatial prior.^49,50^ In neurocomputational terms, this prior may interact with sensory signals by influencing tuning curve distributions,^51^ lateral intraareal connectivity strength,^52^ and/or neural population dynamics.^53^

Another possibility is that the Bayesian computation is more distributed and may include other regions. One likely candidate is the posterior parietal cortex, based on its role in sensorimotor transformations and multisensory integration^54–56^ and its coding and updating of body representation.^57,58^ It has been proposed that posterior parietal areas estimate the state of the body relative to the environment.^59^ Future research employing electrophysiological measures is needed to determine the temporal dynamics and neural implementation of the Bayesian computations within the somatosensory processing hierarchy.

We conclude that perceptual biases are driven by near-optimal somatosensory computations, rather than a distorted hand model. This new perspective on the nature of somatosensory computations allows for a reinterpretation of the ‘hand paradox’. That is, given that the likelihoods and priors underlying hand perception accurately reflect hand geometry, there appears to be no conflict with optimal motor control. The relationship between hand perception and manual actions^4,60^ is an open question, given that manual control relies on the dynamic predictions of a forward model and not a static prior. However, regardless of this relationship, within the Bayesian framework all computations are optimal and the geometry of all representations is unbiased. This novel proposal opens a new window on the understanding of body representations.

## Supplemental Information

### Computational modeling

#### Bayesian model of finger spatial perception

To model the kinematics of a system such as the finger, it is useful to consider it as a kinematic chain, thereby specifying the relevant degrees of freedom. For simplicity, we assume that the finger only flexes or extends, as realized by a combination of one-axis rotations of the three interphalangeal joints (i.e., metacarpophalangeal joint (MC), proximal interphalangeal joint (PIP) and distal interphalangeal joint (DIP)). If the hand is fixed in space, the locations of PIP, DIP, and fingertip (FT), as probed in the experiment, can be expressed in a 2D Cartesian frame of reference, centered at MC, with the y-axis such that it is aligned with the finger when the finger is fully extended, and the x-axis orthogonal to it (see **Figure 1A**).

We define the rotation angles of MC, DIP and PIP, representing flexion/extension motion, as *θ*_1_, *θ*_2_ and *θ*_3_, and the lengths of the base, middle, and tip phalanges as *L*_1_, *L*_2_, and *L*_3_, respectively. Assuming these variables as noiseless, the spatial (Cartesian) location of PIP (***l***_1_), DIP (***l***_2_) and FT (***l***_3_), defined as 2D vectors, ***l***_*n*_ = *f*(*θ*_1_, …, *θ*_*n*_., *L*_1_, …, *L*_*n*_), are geometrically specified as follows:

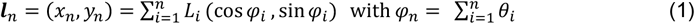

Because noise is ubiquitous in the nervous system, a neural computation of Eq. (1) relies on noisy sensory inputs of the involved joint angles and phalanx lengths, yielding outcomes that are described not as single points but rather as probability distributions over possible positions, as formalized by Bayes’ theorem.

To provide a theoretical framework that explains the observed responses, we designed a probabilistic model of the kinematic chain that assumes optimal Bayesian processing of all potentially relevant signals. For each ***l***_*n*_, we specify the likelihood and the prior with means ***μ***_*lik*_ and ***μ***_*pr*_, respectively, and diagonal covariance matrix *∑*_*lik*_ and *∑*_*pri*_. Strictly speaking, the likelihood follows from the sensory measurements, which we assume to be unbiased but contaminated with Gaussian noise. This is referred to as the measurement distribution, e.g., the distribution of sensory signals for a particular joint angle. However, the brain must perform the inverse approach to compute the sensory likelihood, e.g., the joint angle that has caused the sensory signal that it receives. If the measurement distribution is Gaussian with homogenous variance, the likelihood will be Gaussian with the same variance but centered on the noisy measurement in the specific trial.

Following Bayes’ rule, the posterior is obtained from the product of the likelihood and prior. In matrixform, this is formalized as follows:

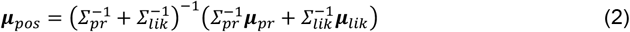

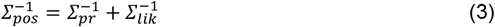

where Eq. (2) is the posterior mean and Eq. (3) is the posterior variance-covariance matrix. Note that Eq. (3) represents *∑*_*pos*_ on a single trial, while experimentally, response variability (*∑*_*r*_) is measured across trials. The response distribution has the same mean ***μ*** of the posterior in a single trial (Eq. 2), but its covariance follows from:^19,20^

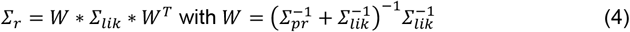

We consider two distinct model variants that formalize how the sensory-to-space transformation (Eq. 1) and Bayesian Integration (Eqns. 2 and 3) are implemented in the somatosensory processing hierarchy (**Figure 1B**).

In **model variant 1** (**Figure 1B**, top, **Figure 1C**, blue), Bayesian integration occurs *before* the sensory-to-space transformation. That is, the likelihood and the prior in Eqns.(2-3) are defined in sensory coordinates: The likelihood is specified as a Gaussian distribution with its mean ***μ***_*lik*_ defined by the involved rotation angles (*θ*_1_, …, *θ*_*n*_) and phalanx lengths (*L*_1_, …, *L*_*n*_). The likelihood variability *∑*_*lik*_ is defined as a diagonal variance-covariance matrix specifying variance in sensory coordinates for joint angles 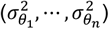 and phalanx lengths 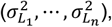 and off-diagonal values set to zero (assuming uncorrelated noise). The prior is also defined in sensory coordinates and specified as Gaussian with mean ***μ***_*pr*_ and a variance-covariance matrix *∑*_*pr*_, allowing covariance between subsequent joint angles (i.e., between *θ*_1_ and *θ*_2_, *θ*_2_ and *θ*_3_).^61,62^ The final step of this model variant transforms the sensory posteriors of the joint angles and segment lengths into proprioceptive localization behavior in space-based coordinates via Eq. (1). A coordinate transformation is therefore necessary. For each ***l***_*n*_, as a linear approximation, we use the Jacobian *J*_*n*_, derived from Eq. (1) by 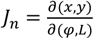, to transform the variance-covariance matrix of the sensory posterior (*∑*_*sens*_) to a variance-covariance matrix in spatial coordinates (*∑*_*spat*_), following:

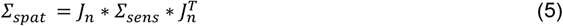

In **model variant 2** (**Figure 1B**, bottom, **Figure 1C**, green), Bayesian integration occurs *after* the sensory-to-space transformation. The likelihood can be initially described by signals related to the individual joint angles and segment lengths of the finger (i.e., sensory coordinates). These sensory signals are first transformed into the Cartesian position of the finger ***l***_*n*_ (via Eqns. 1 and 5), leading to a spatial likelihood with mean ***μ***_*lik*_ and covariance *∑*_*lik*_ defined in Cartesian coordinates. The posterior spatial estimate of finger posture is defined as the product of this spatial likelihood and a spatial prior (via Eqns. 2-3), both represented in Cartesian coordinates.

#### The distorted hand model - the classical hypothesis

We compared the predictions of the Bayesian model to the predictions of the distorted hand model, the classical hypothesis that assumes inbuilt biases in finger representation.^1,6,8^ To this end, we formulated the distorted hand model from Eq. (1), thereby including a constant bias to each joint angle (*θ*_*i*_ + *e*_*θi*_ ) and phalanx length (*e*_*Li*_*L*_*i*_). To make the model probabilistic, for each ***l***_*n*_, we specified a diagonal covariance matrix for joint angles and phalanx lengths, and off-diagonal values set to zero (assuming uncorrelated noise).

### Participants

Twenty healthy volunteers (12 females) provided their informed consent to participate in the experiment. Eighteen participants were included in the analyses (see Data preprocessing). Participants were all right-handed (Edinburgh Handedness Inventory Short Form,^63^ score > 75) and had a mean age of 21.55 ± 2.63 years. They all received course credits as compensation for participating in the experiment. The research was approved by the Ethics Committee of the Faculty of Social Sciences, Radboud University.

**Figure S1.**
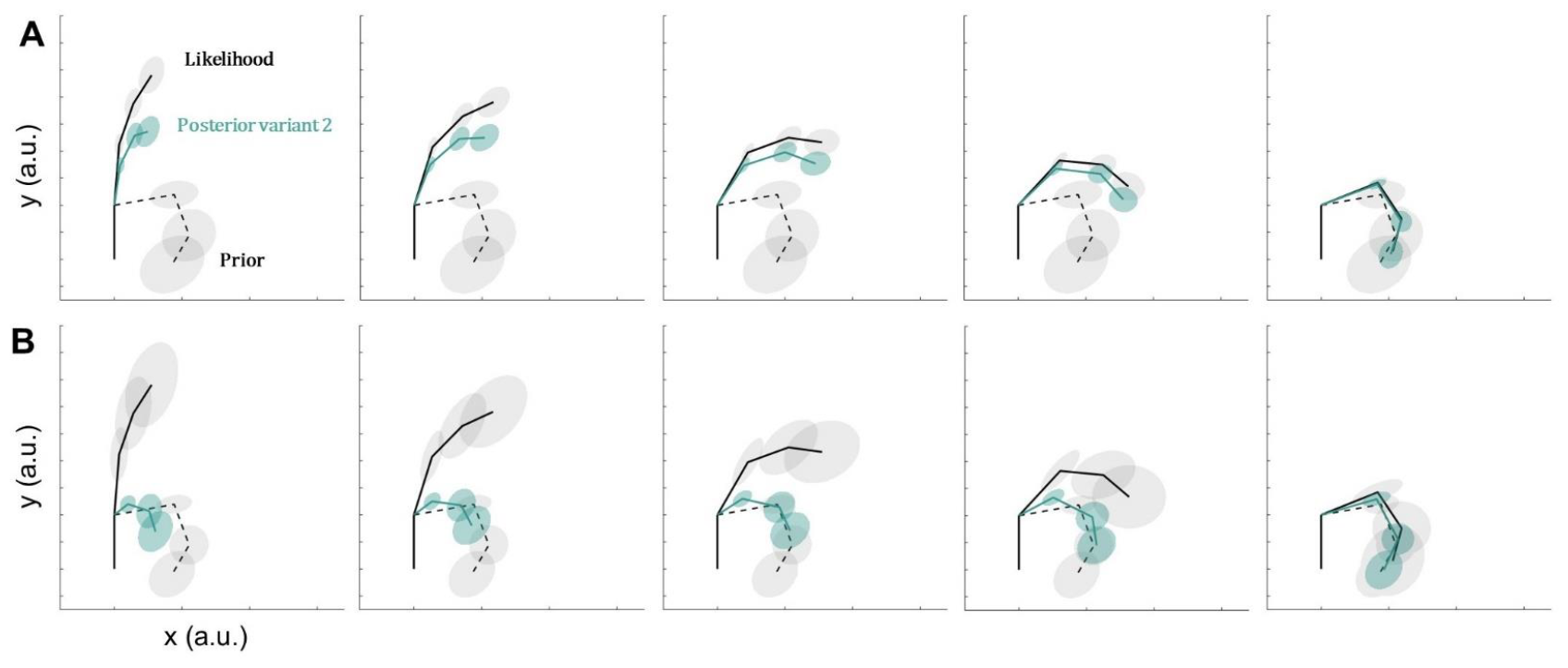
Model simulations: variant 2 predicts posture-dependent biases. **A**. Simulation of spatial posterior for model variant 2 (green) across six different likelihood means, i.e., postures (solid gray). Under a fixed prior (dashed gray), the bias affecting perceived finger geometry is posture dependent. **B**. Biases vary as a function of the relative weight of the likelihood and prior. In bottom panels, the prior’s weight is greater relative to the likelihood, yielding different outcomes in perceived finger configuration and geometry.

### Position tracking and VR

We tracked the position of the index finger via infrared motion tracking (Optotrak Certus, Northern Digital Inc.), with a sampling rate of 100Hz. We used First Principles software to align coordinate systems of all position sensors into a global room-based coordinate system (i.e., registration). We used custom-made rigid bodies for each individual limb segment (forearm, hand dorsum, and the three phalanxes of the index finger) to track the position of the wrist, MC, PIP, DIP, and FT. Time stamps from the VR system were used to segment the continuous 3D signals into blocks corresponding to experimental conditions.

The VR environment was created with Unity. Participants wore an HTC Vive VR headset, while comfortably seated in a chair with their head supported by a chin rest. Images were presented with 1080*1200 pixel resolution to each eye at a 90 Hz refresh rate. The interocular distance was adjusted for each participant such that images were viewed with a field of view of 110°. The VR space was aligned with Optotrak space using a calibration procedure supported by custom-made Python script. The alignment error was within 0.4 and 1.2 mm for all testing sessions.

### Task and procedures

The experiment employed a VR finger mapping task to measure position estimates of the unseen left index finger joints (PIP, DIP) and FT. Participants were instructed to lay their left forearm on the table in front of them. Position estimates were made while holding the left index finger in six unique postures, imposed by asking participants to firmly grasp cylindric objects of three different sizes (40, 60, 80 mm diameter, 200 mm height) in two orientations (**Figure 2A** and **Figure S2, SI**). Each cylinder was used to test two different finger configurations for a total of six unique finger configurations.

The VR environment (**Figure 2B**) presented a dark three-dimensional space with horizon lines, and a realistic screen displaying written instructions. The actual locations of their left index MC and wrist were rendered as blue spheres (diameter 5 mm). This was done to measure perceptual biases related to finger configuration, rather than biases pertaining to the perceived position and configuration of the whole upper limb. Additionally, information on their total index finger length was conveyed via a yellow circumference, centered on the participant’s left MC. The radius of the circle represented participant’s finger length but did not provide any information about the spatial location of targets of interest (PIP, DIP, FT). This experimental choice was merely made to rule out biases in the perception of the virtual environment.

Participants were asked to hold a controller with their right hand, which included a virtual stick and pointer. On each trial, participants started with the virtual pointer at the start position (pointer held in the direction of the virtual monitor). The target of each trial (e.g., ‘fingertip’) was displayed on the virtual monitor. Participants were instructed to indicate with the virtual pointer the perceived location of e.g., their left index fingertip in space, and press a response button to confirm the response. A green wheel appeared on the pointer to indicate that the response was submitted. After each trial participants returned the right arm to the start position.

The experiment included six blocks, each testing one specific finger configuration. Each block contained 100 randomized trials, including 30 trials for PIP, DIP, and FT, and 5 trials for wrist and MC. Trials were self-paced and each block lasted approximately 10 minutes. In total, each participant performed 600 trials. Prior to the finger mapping task, participants completed a practice session to familiarize themselves with the virtual environment and task structure.

**Figure S2.**
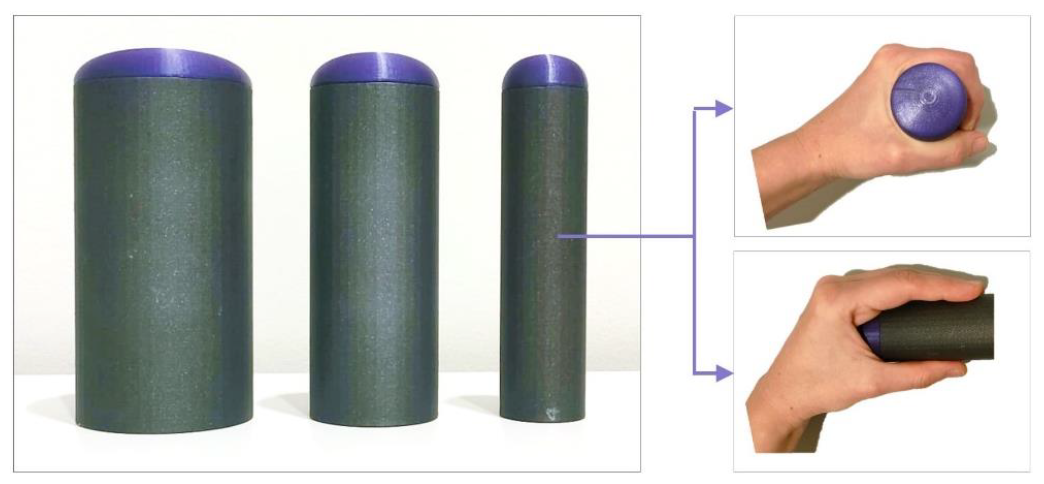
Cylindric volumes. Three cylindric objects (diameters: 40, 60, 80 mm), in two orientations, were used to probe six unique finger postures in each participant. In each block, participants were asked to firmly and stably grasp one of the three objects by keeping the object vertical (e.g., in top-right panel) or horizontal (e.g., in bottom-right panel).

### Data preprocessing

For each participant, motion tracking data and behavioral responses were extracted in 3D coordinates and preprocessed using custom-made MATLAB code (R2022b). Two participants were excluded from the analysis as their data suggested they misunderstood the task instructions. We ascertained participants kept their finger position constant during each block (variation < 1 mm across blocks and participants).

We set out to analyze the data in a 2D plane, i.e., the rotational flexion-extension plane of the finger, as these were the axes upon which finger postures in our task varied, as well as for consistency with theoretical description above. Then, to subtract general biases in localization affecting all positions equally, we aligned the physical and perceived position of the MC to [0,0]. We then rotated each block of responses to align them on the physical MC-wrist axis, to align the perceived and physical hand dorsum and wrist.

For each distribution of estimates (i.e., for each participant, posture, and finger landmark), we aimed to exclude extreme (p<.001) multivariate outliers based on their statistical distance from the geometrical median of the distribution.^64^ This led to the exclusion of on average 9.1% datapoints for each subject (in total: 891 datapoints out of 10800). **Figure S3** (**SI**) shows the data after alignment and outlier removal, with Cartesian positions normalized for total finger length (i.e., [x y]/total finger length).

We calculated the mean physical positions of wrist, MC, PIP, DIP and FT across blocks. These were used to extract fixed parameters for individual model fitting. In detail, we first computed the physical angles (in degrees) between the three vectors defined based on the relative positions of MC-PIP, PIP-DIP and DIP-FT. Then we calculated the length of each phalanx as the Euclidean distance (in mm) between MC-PIP, PIP-DIP, DIP-FT.

**Figure S3.**
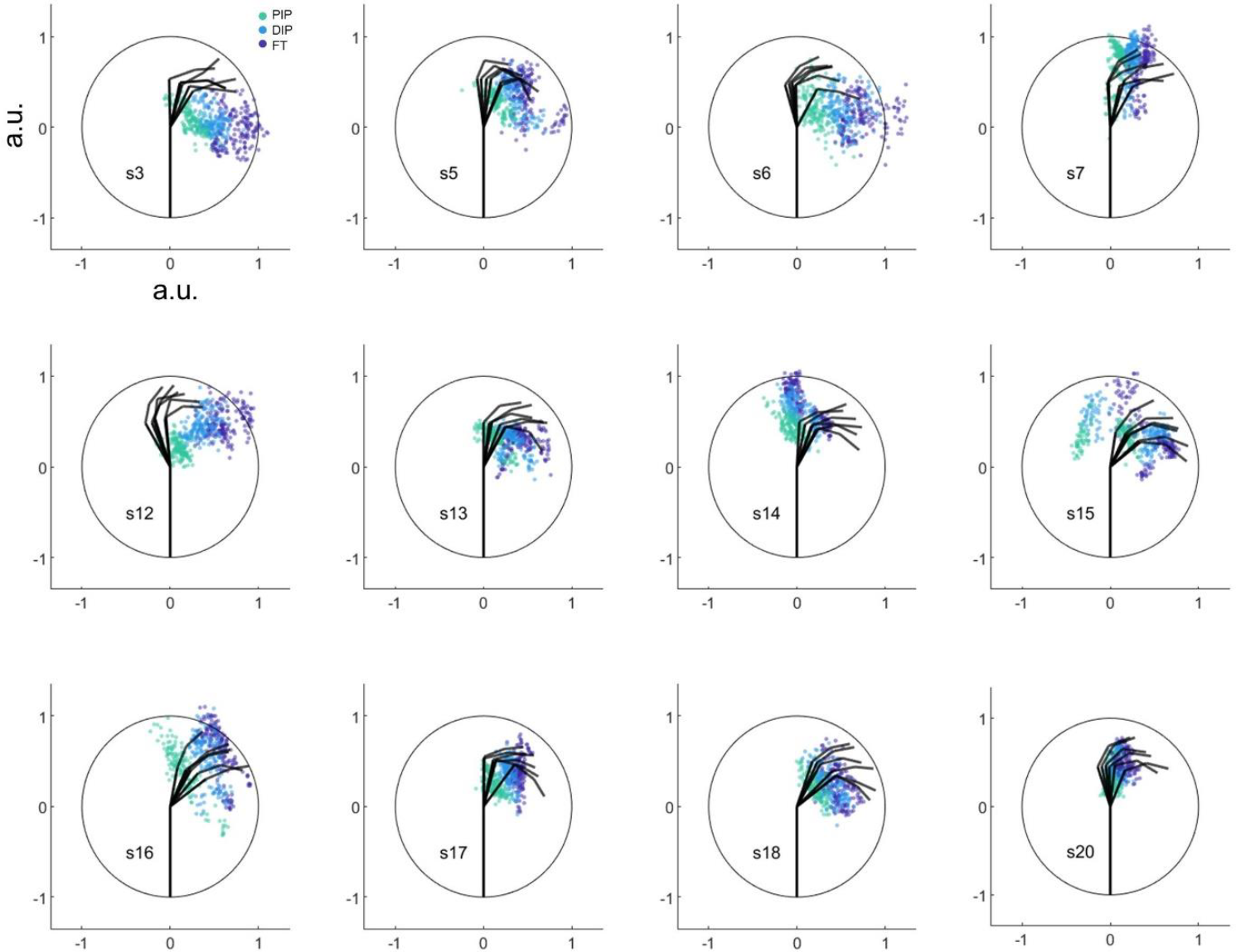
Examples of preprocessed data. Each polar plot shows the normalized circumference (i.e., rendered yellow circle) centered on participant’s MC, along with the six tested postures (black segments), and behavioral responses for PIP (green), DIP (blue), FT (purple).

### Model fitting and analyses

The current section describes how we fit each model to the participants individual responses.

#### Structure of the Bayesian models

For the *likelihood*, we assumed that the sensory measurements (***θ*** and ***L***) were noisy but accurate. The parameter for the mean of each estimate was therefore fixed to its empirical value, i.e., the joint angle on that trial or the actual segment length. The parameter for the variance of each estimate was *free* to vary within realistic constraints. To reduce the number of free parameters, we assumed that likelihood variance over joint angle was similar across joints, and variance over phalanx length was similar across phalanges. We also assumed that the noise in the sensory measurements was uncorrelated. Off-diagonal values for likelihood covariances were therefore fixed at zero. In total, the fitting for the likelihood had six fixed parameters and two free parameters.

For the *prior*, we had different assumptions for signals encoding joint angles and phalanx length. The parameter for the prior means over joint angles was *free* to vary within realistic constraints. In contrast, we assumed accurate priors over phalanx length; we thus fixed the parameter for the means of these priors to their empirical values. The rationale behind these assumptions lies in the biomechanical features of the finger: while joint angles continuously vary, the length of each phalanx is stable in the short-term. It follows that sensory signals on phalanx lengths reflect the actual finger geometry, leading to an accurate measurement and prior distribution. Regarding the prior variance-covariance, we let the variance of each estimate free to vary within realistic constraints. We set the correlation between two sets of joint-pairs as free to vary, given known covariances between them.^61,62^ All other off-diagonal values were set at 0. In total, the fitting for the prior had three fixed parameters and eleven free parameters.

The full model therefore had 22 parameters (13 free, **Table S1, SI**) that were input into the functions described by Eqns. (1-5) via custom-made MATLAB code, following the computational steps of model variant 1 or model variant 2. It is critical to note that the fitting for each model variant had the same free and fixed parameters.

**Table S1.**
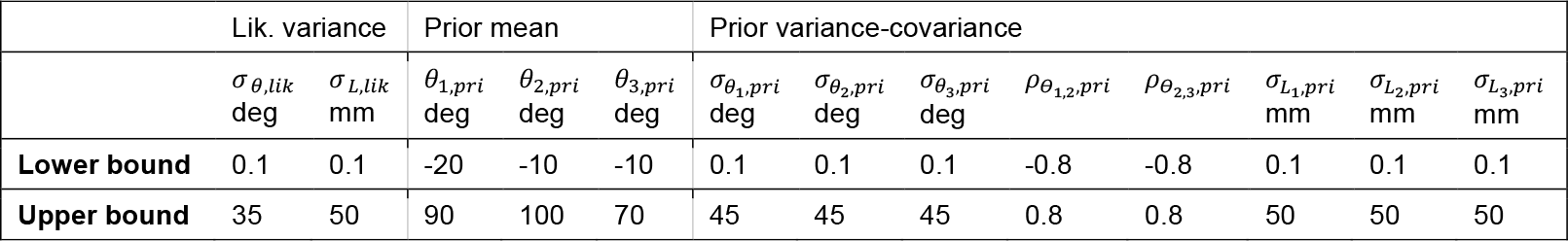
Hard boundaries for model parameter search space.

#### Structure of the distorted hand model

The distorted hand model had six fixed parameters corresponding to true joint angle and phalanx lengths (*θ*_1_, *θ*_2_, *θ*_3_, *L*_1_, *L*_2_, *L*_3_) and 12 free parameters, which correspond to the constant bias on joint angles and phalanx lengths (*e*_*θ*1_, *e*_*θ*2_, *e*_*θ*3_, *e*_*L*1_, *e*_*L*2_, *e*_*L*3_) and the six diagonal values of the variance-co-variance matrix ∑. The lower and upper bounds were [-50, 50] for *e*_*θi*_, [0.2, 1.8] for *e*_*Li*_, [0.1, 35] for diagonal values referring to variance on joint angle, and [0.1, 50] for diagonal values referring to variance on phalanx length.

#### Model fitting and evaluation

To estimate best fitting parameters for each participant and model variant, we maximized the probability of the data (observed behavioral estimates expressed in 2D spatial coordinates), given a range of initial parameters and model variant. We applied Maximum Likelihood Estimation (MLE) to maximize the log-likelihood, with log-likelihood based on a 2D Gaussian function and calculated as:

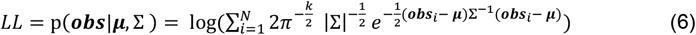

where N is the number of observations; k is the number of dimensions (2); **obs** represent observed localization judgements (expressed in 2D spatial coordinates), relative to MC position (0,0); **μ** and ∑ are the mean and covariance of the response distribution (expressed in 2D spatial coordinates), given each initial set of free parameters. Optimization of the -LL was done using *fmincon* (MATLAB) as done previously.^20,65^ For every model variant and participant, we computed 100 fits using random parameter initializations. We then selected the best fit as the one associated with the lowest -LL (**Table S2, SI**).

For each best fit we then calculated the BIC (Bayesian Information Criterion). This expresses the maximized log-likelihood (LL) as a function of the number of observations (N) and free parameters (k),

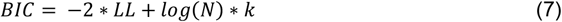

Lower BICs indicate better fits. We calculated the Bayes Factor (BF)^66^ representing the likelihood of model variant 2 relative to model variant 1 (BF_v2,v1_) as 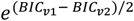 inputting the median difference between *BIC*_*v*1_ and *BIC*_*v*2_.

If model variant 2 well approximates behavioral estimates, we would expect that the observed finger length estimation bias (estimated finger length – actual finger length) / actual finger length *100 varies as a function of the distance between the prior and the likelihood, as shown by model simulations (**Figure 1D**). This distance was quantified as Bhattacharya distance (a variation of the Mahalanobis distance), computed based on the fitted likelihood and prior expressed in Cartesian coordinates:

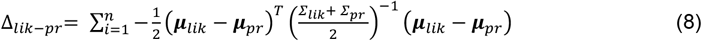

**Table S2.** Fitted parameter values and fit outcome for each subject and model variant.

| Sub<br>ject | Va-<br>riant | Likelihood |  | Prior |  |  |  |  |  |  |  |  |  |  | Fit outcomes |  |  |
| --- | --- | --- | --- | --- | --- | --- | --- | --- | --- | --- | --- | --- | --- | --- | --- | --- | --- |
| | | $\sigma_{\theta,lik}$ | $\sigma_{L,lik}$ | $\theta_{1,pri}$ | $\theta_{2,pri}$ | $\theta_{3,pri}$ | $\sigma_{\theta_{1,pri}}$ | $\sigma_{\theta_{2,pri}}$ | $\sigma_{\theta_{3,pri}}$ | $\sigma_{L_{1,pri}}$ | $\sigma_{L_{2,pri}}$ | $\sigma_{L_{3,pri}}$ | $\rho_{\theta_{1,2,pri}}$ | $\rho_{\theta_{2,3,pri}}$ | -LL | BIC | R<sup>2</sup> |
| 1 | 1 | 16.33 | 39.68 | -20.00 | 100.00 | -10.00 | 35.19 | 17.60 | 31.82 | 29.76 | 17.12 | 26.62 | 0.80 | 0.54 | 4060.30 | 8202.40 | 0.03 |
|  | 2 | 6.69 | 12.67 | -20.00 | -10.00 | 64.43 | 16.45 | 32.90 | 45.00 | 31.64 | 50.00 | 25.00 | -0.75 | -0.80 | 3830.39 | 7742.57 | 0.40 |
| 2 | 1 | 35.00 | 50.00 | 27.01 | 100.00 | 36.40 | 19.89 | 9.95 | 19.89 | 39.52 | 19.76 | 19.19 | 0.67 | -0.80 | 4011.85 | 8105.50 | -0.23 |
|  | 2 | 8.78 | 32.51 | 78.62 | 92.64 | 26.10 | 18.03 | 36.05 | 45.00 | 17.64 | 23.15 | 29.94 | -0.80 | -0.80 | 3704.18 | 7490.15 | 0.41 |
| 3 | 1 | 24.32 | 50.00 | 90.00 | 81.89 | -10.00 | 40.98 | 26.09 | 13.04 | 39.13 | 19.56 | 12.48 | -0.80 | 0.80 | 9233.43 | 18548.64 | -7.01 |
|  | 2 | 24.81 | 37.77 | 90.00 | 59.03 | -10.00 | 25.34 | 33.15 | 45.00 | 27.52 | 29.18 | 20.92 | -0.80 | 0.23 | 4179.01 | 8439.82 | 0.59 |
| 4 | 1 | 16.92 | 50.00 | -20.00 | 5.84 | -10.00 | 30.34 | 45.00 | 40.52 | 35.09 | 17.54 | 31.84 | 0.80 | 0.80 | 6575.50 | 13232.79 | -1.29 |
|  | 2 | 35.00 | 50.00 | -20.00 | -10.00 | 48.11 | 45.00 | 45.00 | 45.00 | 40.58 | 20.29 | 20.23 | -0.80 | -0.80 | 4649.06 | 9379.90 | -0.17 |
| 5 | 1 | 11.25 | 50.00 | 90.00 | 100.00 | -10.00 | 45.00 | 22.50 | 16.07 | 29.02 | 15.33 | 30.65 | -0.37 | 0.80 | 4539.42 | 9160.63 | -0.78 |
|  | 2 | 13.69 | 12.40 | 90.00 | 79.58 | -1.33 | 19.14 | 14.54 | 29.08 | 18.19 | 19.88 | 39.75 | -0.80 | 0.39 | 3695.98 | 7473.75 | 0.67 |
| 6 | 1 | 25.89 | 50.00 | 90.00 | 40.53 | -10.00 | 39.23 | 33.17 | 21.24 | 34.27 | 17.14 | 8.57 | -0.80 | 0.80 | 4738.69 | 9559.18 | -1.04 |
|  | 2 | 22.50 | 31.77 | 87.00 | 28.97 | 32.70 | 37.42 | 45.00 | 45.00 | 18.71 | 14.27 | 18.84 | -0.79 | 0.04 | 3989.29 | 8060.38 | 0.67 |
| 7 | 1 | 35.00 | 50.00 | 8.94 | 35.40 | -10.00 | 12.04 | 23.57 | 45.00 | 50.00 | 25.00 | 18.71 | -0.80 | -0.60 | 3945.41 | 7972.61 | -0.23 |
|  | 2 | 35.00 | 45.43 | 6.06 | 36.57 | -4.24 | 15.50 | 31.00 | 15.50 | 50.00 | 25.00 | 14.94 | -0.80 | 0.32 | 3922.63 | 7927.04 | -0.27 |
| 9 | 1 | 32.56 | 50.00 | -9.22 | 100.00 | 70.00 | 32.59 | 45.00 | 26.29 | 34.05 | 17.03 | 8.51 | -0.80 | 0.80 | 4248.01 | 8577.82 | 0.69 |
|  | 2 | 35.00 | 50.00 | 7.29 | 49.58 | 49.38 | 34.00 | 45.00 | 22.50 | 34.70 | 22.29 | 11.14 | -0.64 | -0.34 | 4253.61 | 8589.02 | 0.52 |
| 10 | 1 | 4.49 | 41.26 | -19.80 | 99.79 | -9.66 | 13.41 | 18.55 | 37.10 | 46.33 | 29.70 | 35.43 | 0.80 | 0.72 | 7278.72 | 14639.23 | 0.07 |
|  | 2 | 27.65 | 50.00 | -20.00 | 58.17 | 50.80 | 42.92 | 45.00 | 45.00 | 49.76 | 39.09 | 19.55 | -0.80 | -0.80 | 4542.96 | 9167.71 | 0.20 |
| 12 | 1 | 35.00 | 50.00 | 24.32 | 71.15 | -10.00 | 18.53 | 37.07 | 44.91 | 44.81 | 22.40 | 25.37 | -0.80 | -0.61 | 4200.77 | 8483.33 | 0.29 |
|  | 2 | 18.35 | 50.00 | 59.11 | -10.00 | 30.20 | 23.93 | 22.50 | 45.00 | 26.54 | 15.96 | 20.88 | -0.63 | -0.79 | 3918.52 | 7918.82 | 0.72 |
| 13 | 1 | 18.63 | 50.00 | -13.90 | 100.00 | 70.00 | 26.43 | 13.21 | 26.43 | 38.01 | 23.87 | 11.93 | -0.63 | -0.68 | 4200.63 | 8483.04 | 0.08 |
|  | 2 | 12.13 | 16.98 | 90.00 | 100.00 | 70.00 | 22.84 | 45.00 | 45.00 | 32.22 | 16.11 | 8.05 | -0.80 | -0.48 | 4085.31 | 8252.41 | 0.39 |
| 14 | 1 | 12.36 | 50.00 | 68.03 | 2.55 | 70.00 | 40.27 | 45.00 | 22.50 | 35.66 | 17.83 | 33.88 | -0.70 | 0.80 | 7303.49 | 14688.76 | -5.47 |
|  | 2 | 22.50 | 16.20 | -20.00 | -10.00 | 21.93 | 17.02 | 33.89 | 45.00 | 34.75 | 26.95 | 13.47 | -0.48 | -0.80 | 4088.19 | 8258.17 | 0.24 |
| 15 | 1 | 35.00 | 50.00 | -20.00 | 68.60 | 49.91 | 45.00 | 22.50 | 23.46 | 26.11 | 31.28 | 15.64 | 0.80 | -0.53 | 4285.28 | 8652.35 | 0.40 |
|  | 2 | 35.00 | 25.52 | -20.00 | 98.18 | 54.15 | 45.00 | 22.50 | 25.57 | 22.76 | 29.08 | 22.77 | 0.64 | -0.80 | 4229.53 | 8540.84 | 0.44 |
| 16 | 1 | 20.24 | 50.00 | -19.07 | 100.00 | -10.00 | 45.00 | 23.44 | 19.33 | 30.89 | 15.45 | 7.72 | 0.21 | 0.80 | 14149.87 | 28381.54 | -13.20 |
|  | 2 | 31.25 | 12.92 | 3.04 | 100.00 | 21.37 | 31.11 | 25.90 | 38.09 | 42.37 | 21.19 | 10.59 | 0.80 | -0.80 | 4028.99 | 8139.77 | 0.56 |
| 17 | 1 | 35.00 | 50.00 | 12.33 | 100.00 | 50.37 | 20.95 | 19.15 | 22.40 | 39.83 | 19.92 | 9.96 | 0.43 | -0.80 | 4074.64 | 8231.07 | -0.48 |
|  | 2 | 9.05 | 22.58 | 90.00 | 60.37 | 16.38 | 19.33 | 38.66 | 45.00 | 23.45 | 46.91 | 50.00 | -0.80 | -0.80 | 3994.62 | 8071.02 | 0.18 |
| 18 | 1 | 20.69 | 50.00 | -8.85 | 100.00 | 70.00 | 30.05 | 15.03 | 30.05 | 36.69 | 18.34 | 23.74 | -0.12 | -0.80 | 4104.04 | 8289.88 | -0.37 |
|  | 2 | 10.45 | 32.93 | 90.00 | 57.42 | 28.66 | 22.35 | 44.69 | 45.00 | 24.31 | 40.81 | 36.47 | -0.80 | -0.80 | 3917.11 | 7916.00 | 0.34 |
| 19 | 1 | 35.00 | 31.31 | -20.00 | 100.00 | 62.26 | 34.24 | 17.12 | 34.24 | 45.82 | 30.33 | 26.79 | 0.16 | -0.80 | 4405.51 | 8892.80 | -0.11 |
|  | 2 | 32.45 | 50.00 | -20.00 | 100.00 | 46.55 | 38.31 | 45.00 | 45.00 | 42.54 | 21.27 | 42.54 | 0.42 | -0.61 | 4344.07 | 8769.94 | -0.26 |
| 20 | 1 | 13.52 | 50.00 | 23.92 | -10.00 | 18.41 | 10.12 | 7.60 | 3.80 | 50.00 | 50.00 | 25.00 | -0.73 | -0.14 | 4083.99 | 8249.76 | -4.15 |
|  | 2 | 11.16 | 20.44 | 8.80 | -10.00 | 14.81 | 10.35 | 20.70 | 41.39 | 50.00 | 50.00 | 25.00 | -0.79 | -0.52 | 3884.20 | 7850.19 | -0.60 |
| Mdn | 1 | 22.50 | 50.00 | 0.04 | 99.89 | 4.37 | 31.46 | 22.50 | 24.87 | 37.35 | 19.66 | 21.46 | -0.25 | 0.20 | 4266.65 | 8615.09 | -0.30 |
| IQR | 1 | 18.52 | 0.00 | 45.95 | 52.45 | 69.29 | 19.85 | 14.16 | 13.41 | 9.46 | 7.48 | 14.68 | 1.39 | 1.47 | 2027.30 | 4054.60 | 1.29 |
| Mdn | 2 | 22.50 | 32.14 | 8.04 | 57.79 | 29.43 | 23.38 | 34.97 | 45.00 | 31.93 | 24.07 | 20.90 | -0.80 | -0.79 | 4011.80 | 8105.40 | 0.39 |
| IQR | 2 | 20.75 | 31.02 | 109.25 | 89.63 | 31.44 | 18.26 | 17.82 | 6.08 | 18.26 | 16.10 | 12.80 | 0.17 | 0.42 | 299.44 | 598.88 | 0.36 |

**Figure S4.**
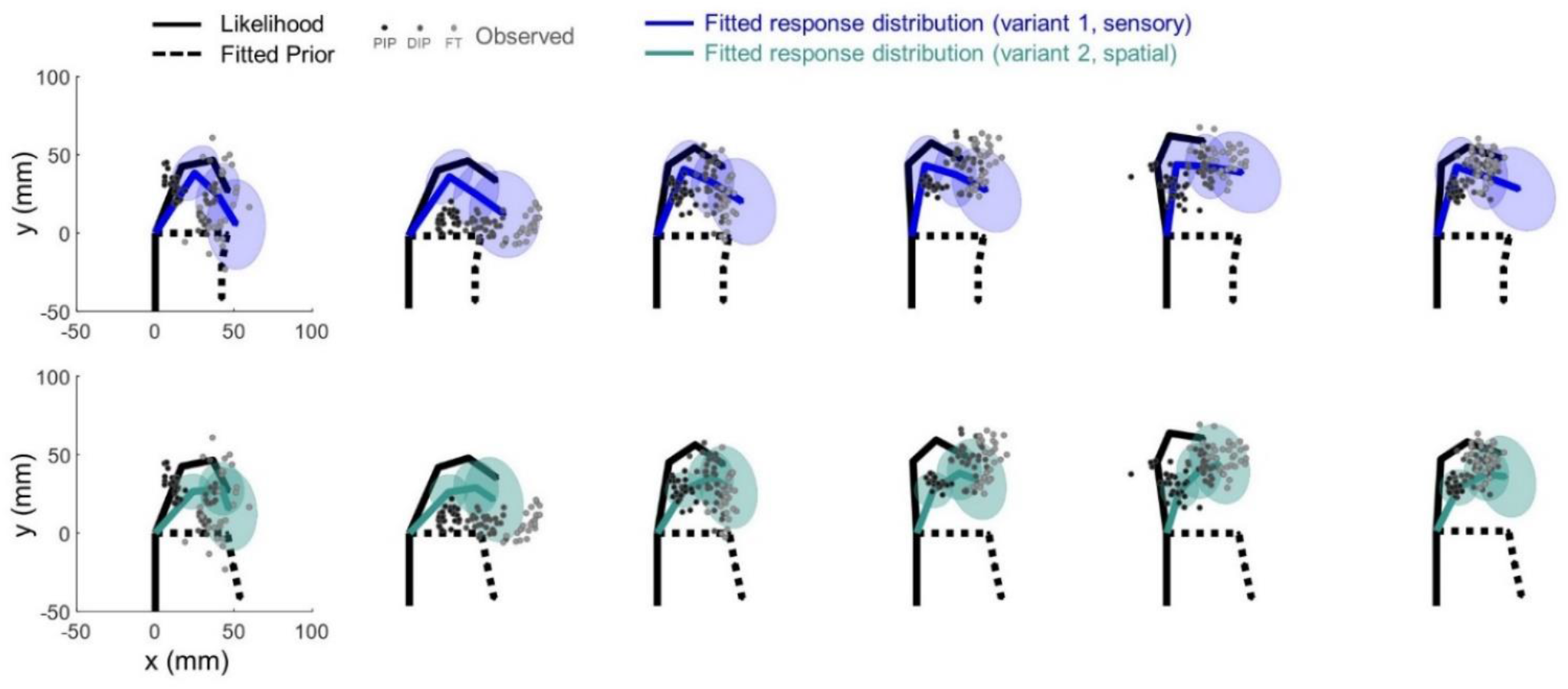
Example of fitting results across postures. Fitting results for one representative participant (variant 1, top panel; variant 2, bottom panel) are plotted in Cartesian coordinates for all the six postures.

### Additional analyses

#### Ruling out truncation/attraction biases

During the task, participants in VR could see the position of their MC and a yellow circumference centered on the MC. The circumference’s radius represented the total length of participant’s index finger when stretched (**Figure 2**). Because the participant’s actual finger was flexed freely within the plane *away from* the circle, none of the visual cues gave information about the spatial location of targets of interest (PIP, DIP, FT).

However, it is still possible that providing this information affected participant’s responses in such a way as to corrupt our estimate of their finger perception. The circle could serve as an anchor, attracting participant’s responses towards it. Alternatively, the circle could serve as a boundary, repelling participant’s responses away from it; this could especially be the case when participant’s fingertips happened to be very close to or overlapping with the circumference (i.e., extended finger posture). We took three approaches to rule out these possibilities.

First, on a theoretical level, if the circle attracted the responses of participants, we would observe an *overestimation* in the measured perception of finger length. This is because responses would be drawn away from the MC (our origin) and towards the circle. Instead, we very clearly observed an underestimation, ruling out attraction as a possibility.

Second, if our results were due to truncation of the fingertip response distribution, we would have expected to observe a greater underestimation of the tip phalanx (i.e., distance between DIP and FT) than the other segments, i.e., base phalanx (MC-PIP) and middle phalanx (PIP-DIP). Indeed, we may not expect to see an underestimation for the latter two segments at all. Instead, as it is visible in **Figure S3** (**SI**), spatial biases affect not only FT, but also DIP and PIP estimation responses. Specifically, segment length misestimation across postures and participants was on average (± sem) -8.65% ± 5.06 for the base segment, -24.68% ± 6.28 for the middle segment, and 1.26% ± 7.82 for the tip segment. One-tailed t-test against zero showed that underestimation is significant for the middle segment (t(17) = -3.93, p = .001), marginally significant for the base segment (t(17) = -1.71, p = .053 and not significant for the tip segment (t(17) = 0.16, p = .563). This pattern of results—especially the lack of underestimation for the tip phalanx—would not be expected if our results were simply due to response truncation.

Third, if our results were solely due to the avoidance of the imposed boundary, we would expect that participants truncated their response distributions at that boundary. In contrast, we consistently observed estimates beyond this boundary. In the polar plots of **Figure S5** (**SI**), the radius of circumferences represents the distance from MC and FT when the finger is extended, i.e., total finger length. The black segments represent the six tested postures, each associated with a different color. Response distributions for the FT are colored accordingly. For most postures and participants fingertip position did not lie close to the circumference. If truncation occurred, we would have expected fingertip estimation responses to cluster *within* the circle. Rather, in some cases (e.g., in subjects 4, 6, 14), responses cross the circle boundary, coherently with non-truncated distributions.

**Figure S5.**
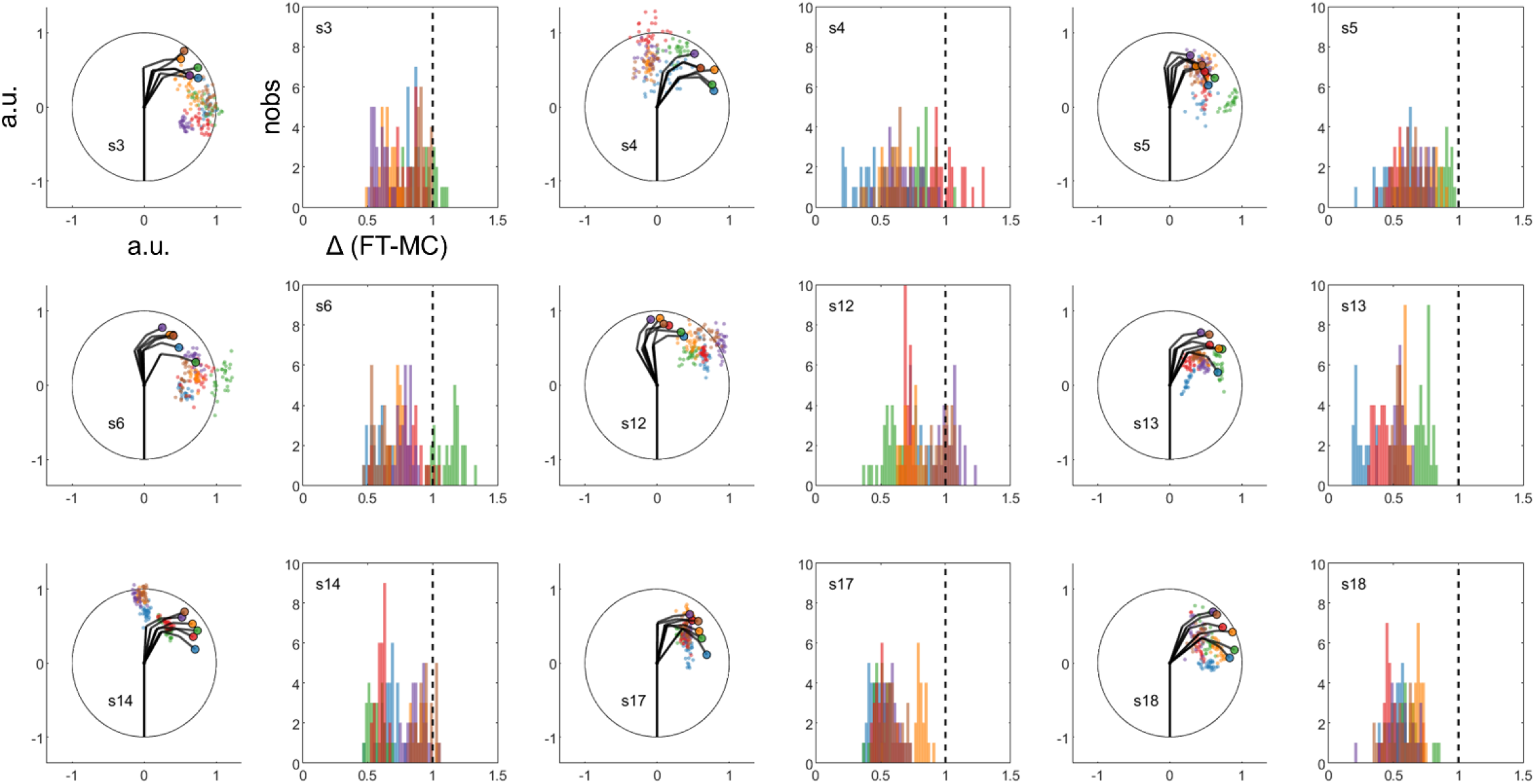
Ruling out truncation/attraction biases. Each polar plot shows the normalized circumference (i.e., rendered yellow circle) centered on participant’s MC, along with the six tested postures (black segments). Fingertip localization responses for a participant (i.e., Cartesian positions normalized on total finger length) are represented as small dots; each color represents a posture. Accordingly, real fingertip positions across postures are plotted as big colored dots. The same colors are used to represent frequency distributions of the estimated distance between MC and FT. Black dashed line represents total finger length.

## Fundings and acknowledgements

V.C.P is supported by the Radboud Excellence Initiative grant. L.E.M is supported by ERC 101076991 SOMATOGPS grant. W.P.M. is supported by the following grants: NWA-ORC-1292.19.298, NWO-SGW-406.21.GO.009 and Interreg NWE-RE:HOME.

We extend our gratitude to the Sensorimotor Lab members for the numerous scientific discussions, and the Technical Support Group of DCC for their invaluable assistance and expertise.

## Data availability

Raw data are available at: <u>di.dcc.DSC_2023.00011_285</u>

Further information and requests for data and code should be directed to the lead contact, Valeria Peviani.

## Author contributions

V.C.P., L.E.M., W.P.M., Conceptualization; Methodology

V.C.P., Funding Acquisition, Investigation, Writing – Original Draft

L.E.M., W.P.M., Resources, Supervision, Writing – Review & Editing

